# The overlapping genetic architecture of psychiatric disorders and cortical brain structure

**DOI:** 10.1101/2023.10.05.561040

**Authors:** Zhiqiang Sha, Varun Warrier, Richard A.I. Bethlehem, Laura M. Schultz, Alison Merikangas, Kevin Y. Sun, Ruben C. Gur, Raquel E. Gur, Russell T. Shinohara, Jakob Seidlitz, Laura Almasy, Ole A. Andreassen, Aaron F. Alexander-Bloch

**Affiliations:** Department of Psychiatry, University of Pennsylvania, Philadelphia, PA, USA; Department of Child and Adolescent Psychiatry and Behavioral Science, The Children’s Hospital of Philadelphia, Philadelphia, PA, USA; Department of Psychiatry, University of Cambridge, Cambridge, UK; Department of Psychology, University of Cambridge, Cambridge, UK; Department of Biomedical and Health Informatics, Children’s Hospital of Philadelphia, Philadelphia, PA, USA; Lifespan Brain Institute, The Children’s Hospital of Philadelphia and Penn Medicine, Philadelphia, PA, USA; Department of Genetics, Perelman School of Medicine, University of Pennsylvania, Philadelphia, PA, USA; Department of Neurology, Perelman School of Medicine, University of Pennsylvania, Philadelphia, PA, 19104, USA; Department of Radiology, Perelman School of Medicine, University of Pennsylvania, Pennsylvania, PA, 19104, USA; Penn Statistics in Imaging and Visualization Endeavor (PennSIVE), Department of Biostatistics, Epidemiology, and Informatics, Perelman School of Medicine, University of Pennsylvania, 423 Guardian Dr, Philadelphia, PA 19104, United States; Center for Biomedical Image Computing and Analytics (CBICA), Department of Radiology, Perelman School of Medicine, United States; NORMENT Centre, Division of Mental Health and Addiction, Oslo University Hospital & Institute of Clinical Medicine, University of Oslo, Oslo, Norway

**Author notes:** **Corresponding author:** Zhiqiang Sha Ph.D., Lifespan Brain Institute at University of Pennsylvania and Children’s Hospital of Philadelphia Department of Child and Adolescent Psychiatry and Behavioral Science, The Children’s Hospital of Philadelphia, Department of Psychiatry, University of Pennsylvania, Aaron Alexander-Bloch, M.D. Ph.D., Lifespan Brain Institute at University of Pennsylvania and Children’s Hospital of Philadelphia Department of Child and Adolescent Psychiatry and Behavioral Science, The Children’s Hospital of Philadelphia, Department of Psychiatry, University of Pennsylvania.

**Keywords:** brain structure, psychiatric disorders, genetic overlap, polygenic risk, transcriptomic profiles

## Abstract

Both psychiatric vulnerability and cortical structure are shaped by the cumulative effect of common genetic variants across the genome. However, the shared genetic underpinnings between psychiatric disorders and brain structural phenotypes, such as thickness and surface area of the cerebral cortex, remains elusive. In this study, we employed pleiotropy-informed conjunctional false discovery rate analysis to investigate shared loci across genome-wide association scans of regional cortical thickness, surface area, and seven psychiatric disorders in approximately 700,000 individuals of European ancestry. Aggregating regional measures, we identified 50 genetic loci shared between psychiatric disorders and surface area, as well as 26 genetic loci shared with cortical thickness. Risk alleles exhibited bidirectional effects on both cortical thickness and surface area, such that some risk alleles for each disorder increased regional brain size while other risk alleles decreased regional brain size. Due to bidirectional effects, in many cases we observed extensive pleiotropy between an imaging phenotype and a psychiatric disorder even in the absence of a significant genetic correlation between them. The impact of genetic risk for psychiatric disorders on regional brain structure did exhibit a consistent pattern across highly comorbid psychiatric disorders, with 80% of the genetic loci shared across multiple disorders displaying consistent directions of effect. Cortical patterning of genetic overlap revealed a hierarchical genetic architecture, with the association cortex and sensorimotor cortex representing two extremes of shared genetic influence on psychiatric disorders and brain structural variation. Integrating multi-scale functional annotations and transcriptomic profiles, we observed that shared genetic loci were enriched in active genomic regions, converged on neurobiological and metabolic pathways, and showed differential expression in postmortem brain tissue from individuals with psychiatric disorders. Cumulatively, these findings provide a significant advance in our understanding of the overlapping polygenic architecture between psychopathology and cortical brain structure.

## Introduction

Psychiatric disorders are widely recognized as brain-related conditions that have a significant genetic component^1–4^. Extensive large-scale genetic analyses utilizing case-control designs have revealed that common genetic variants across the human genome can account for at least 10% of the variability observed in mental disorders^2,5,6^. Moreover, comprehensive mega-analyses encompassing diverse cohorts of individuals with psychiatric disorders have demonstrated that a majority of these disorders are accompanied by alterations in regional brain anatomy, many of which are shared across psychiatric disorders that have high co-morbidity with one another^7–14^. Importantly, these structural changes in the brain have been observed not only in individuals diagnosed with psychiatric disorders but also in their unaffected relatives^15–17^. Moreover, individuals at a higher genetic risk for mental disorders have been reported to display distinct brain structural profiles and exhibit developmental trajectories that deviate from typical neurodevelopment when compared to those at a lower genetic risk^18–20^. However, the neurogenetic mechanisms that underpin the shared genetic architecture between psychopathology and brain structures remain unclear.

Genome-wide association studies (GWAS) conducted on brain MRI anatomical measures in non-clinical samples have consistently revealed significant heritability across nearly all regional brain measures^21,22^. Of these measures, regional surface area (SA), which quantifies the outer area of the cerebral cortex, typically exhibits higher heritability compared to regional cortical thickness (CT), which represents the distance between the outer and inner surfaces of cortical gray matter^22,23^. The two components that determine cortical volume, CT and SA, are not significantly genetically correlated, consistent with distinct cellular mechanisms underlying interindividual variability in these phenotypes^24^. Moreover, only a small number of genetic loci have been implicated in both GWAS of brain structural measures and GWAS of psychiatric disorders^21,22^, suggesting limited pleiotropic effects where a single gene influences both phenotypes. Although psychiatric disorders and brain structures each have strong genetic influences, and GWAS can be leveraged to uncover genetic correlations among different phenotypes^25^, current evidence supporting substantial genetic correlation at the genome level between psychiatric disorders and brain structures remains limited.

The dearth of evidence for widespread shared genetic influences on psychiatric disorders and brain imaging measures may result, in part, because previous studies do not fully leverage information contained in existing GWAS of the respective phenotypes. Some previous studies focus simply on the subset of SNPs that surpass a Bonferroni-corrected threshold of significance in each trait^26,27^. Alternatively, methods such as linkage disequilibrium score regression (LDSC)^28,29^ examine the global genetic correlation between effect sizes of genetic variants associated with two phenotypes^6,22^. However, these approaches may underestimate genetic overlap in the context of a mixture of genetic variants with concordant and discordant directions of effect for two phenotypes. Polygenic scores, which quantify an individual’s genetic predisposition to a specific phenotype by combining the genetic effects of multiple common variants derived from GWAS results^30–32^, have also been used to assess shared genetic influences. For instance, studies utilizing polygenic scoring have reported that certain brain imaging phenotypes are correlated with polygenic risks for psychiatric disorders^21,33,34^, but this approach will likewise underestimate shared genetic influences in the context of a mixture of genetic variants with concordant and discordant directions of effect. Many previous psychiatric imaging-genetic studies have also been relatively underpowered, given the small effect sizes of common genetic variants on phenotypic variability. Collectively, it remains a challenge to comprehensively and reliably capture genetic overlap between traits with complex, polygenic architecture using traditional analytical methods.

Consequently, significant efforts have been devoted to investigating genetic loci and genetic correlations associated with complex phenotypes in reliable, comprehensive, and cost-effective methods that more fully leverage existing GWAS data. Recently, a pleiotropy-informed method called the conjunctional false discovery rate (FDR) has been proposed to enhance the discovery of shared genetic loci between pairs of phenotypes by combining two GWAS summary statistics^35–37^. This Bayesian-based approach determines the extent of genetic overlap between two phenotypes by leveraging the pleiotropic enrichment of genetic variants associated with each phenotype that exhibit increased association with the paired phenotype. One of the advantages of this approach is its ability to detect shared genetic loci with concordant or discordant effect directions between the two phenotypes, e.g. genetic variants associated with psychiatric risk could be associated with greater and/or lesser CT/SA depending on the specific genetic locus, a critical point of distinction that may be obscured in the global genetic correlation^36^. Conjunctional FDR has been successfully employed to identify overlapping genetic associations between psychiatric disorders and neurological disorders, as well as with physical health phenotypes^38–40^. The prior literature suggests the suitability of this method to examine the shared genetic architecture between psychiatric disorders and brain structure.

Here, in order to advance our understanding of genetic underpinnings underlying the relationship between psychiatric disorders and structural brain imaging measurements, we integrated GWAS summary statistics from approximately 700,000 individuals of European ancestry. Psychiatric GWAS data were obtained from the Psychiatric Genomics Consortium, including samples for attention-deficit/hyperactivity disorder (ADHD, N=53,293)^41^, anxiety disorder (ANX, N=17,310)^42^, autism spectrum disorder (ASD, N=46,350)^43^, bipolar disorder (BD, N=51,710)^44^, major depressive disorder (MDD, N=143,265)^45^, post-traumatic stress disorder (PTSD, N=214,408)^46^, and schizophrenia (SCZ, N=130,644)^47^. We conducted independent GWAS analyses for 180 bilaterally averaged CT and SA measures defined by the Human Connectome Parcellation^48^, using data from the UK Biobank and the ABCD study as recently reported (combined N=36,663)^23^. We confirmed that these psychiatric GWAS did not include UK Biobank and ABCD cohorts to eliminate false positive estimates arising from overlapping samples. Psychiatric and neuroimaging GWAS were then integrated in a series of conjunctional FDR analyses. Using this approach, we were able to directly characterize psychiatric risk alleles with both concordant and discordant directions of effect on brain structure. We also differentiated shared risk alleles, and the corresponding pattern of regional CT and SA influenced by these genetic loci, in cases where genetic variants were jointly associated with multiple disorders and in cases where genetic variants were uniquely associated with a single disorder. Finally, we performed functional annotation of shared genetic variants in terms of regulation of gene expression, cell-type specificity, chromatin regulation, and overlap with genes that are differentially expressed in postmortem brain tissue of individuals with psychiatric disorders. Collectively, our results represent a major advance in our understanding of shared genetic influences on psychiatric vulnerability and regional structural variation in the human brain.

## Results

### Genetic overlap between psychiatric disorders and brain structures

We found compelling evidence for polygenic overlap between psychiatric disorders and measurements of CT and SA derived from brain MRI. For each of the 7 neuropsychiatric disorders, there was enrichment between SNPs associated with the psychiatric disorders and SNPs associated with 180 regional CT measurements and 180 regional SA measurements, as assessed by pleiotropic enrichment plots^37^ (Supplementary Figures 1-14). We then conducted conjunctional FDR analysis using the pleioFDR toolbox^37^ to identify shared genetic loci for each specific disorder-brain pair. The conjunctional FDR score for each SNP is defined as the maximum of conditional *FDR(disorder | brain)* and conditional *FDR(brain | disorder)*. For each direction, the conditional FDR score for each SNP is computed by re-ranking the test statistics of the primary phenotype at the genome level by conditioning them on the association with the auxiliary phenotype^37^ (see Methods). Both CT and SA had joint associations with psychiatric disorders at multiple specific genetic loci (Figure 1).

**Figure 1.**
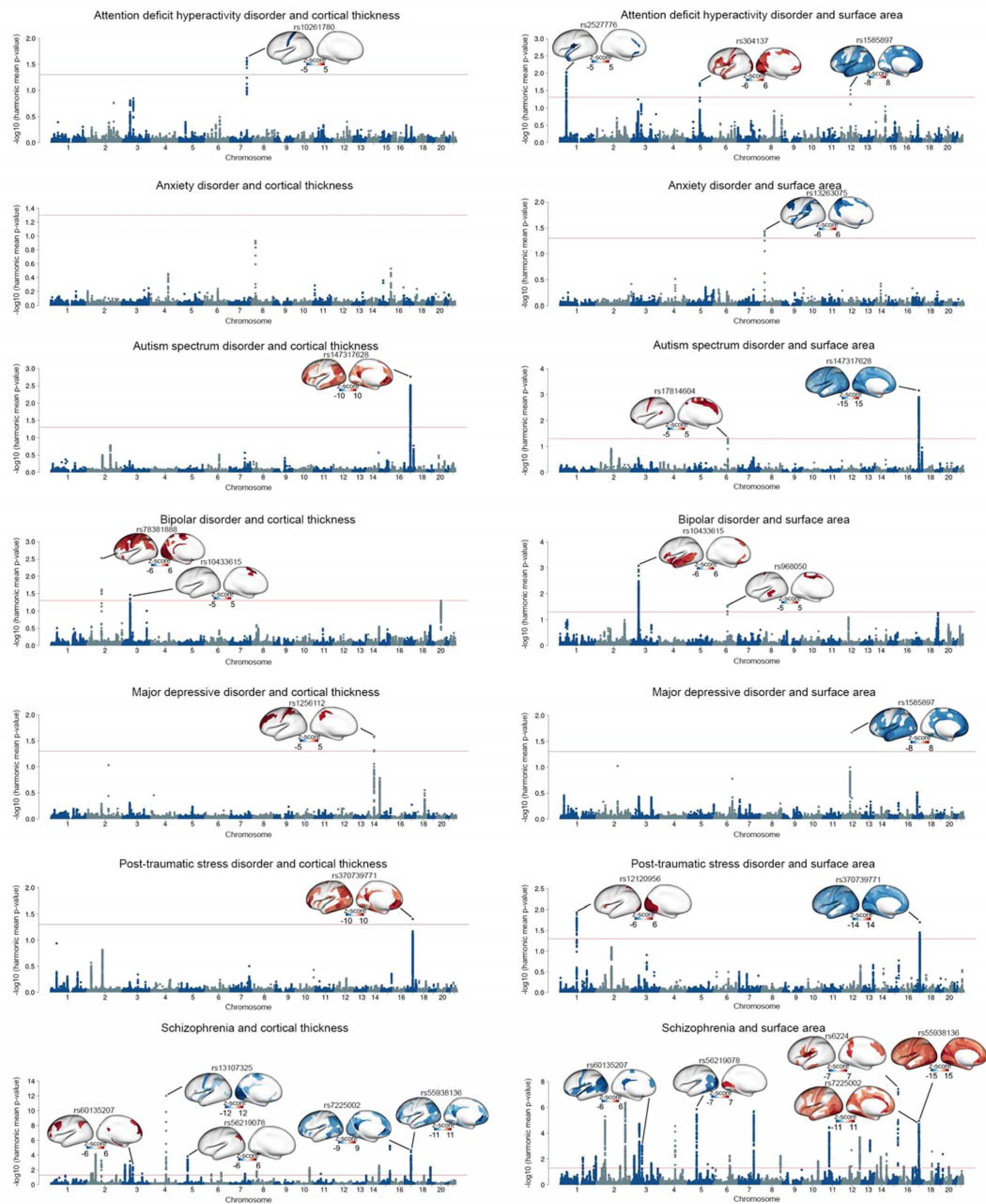
Shared genetic loci between psychiatric disorders and cortical thickness and surface area measures. Manhattan plots illustrating the global conjunctional FDR analyses, highlighting the shared genetic loci between each psychiatric disorder and measures of cortical thickness and surface area. The significance threshold of harmonic mean FDR-corrected p-value<0.05 is represented by red lines. The cortical map illustrates the brain regions that are shared by the lead SNP with the specific psychiatric disorder. Red indicates concordant directions of effect, where greater psychiatric vulnerability is associated with thicker cortex or larger surface area. Conversely, blue represents discordant directions of effect, where greater psychiatric vulnerability is associated with thinner cortex or smaller surface area. Color bars indicate z-scores derived from the conjunctional FDR analysis between psychiatric disorders and brain structure.

Given the large multiple testing burden, we first identified aggregate genome-wide-significant associations between CT or SA and each disorder, using the harmonic mean FDR-corrected p-value (HMP) across brain regions^49^. Using FUMA^50^ to identify independent lead SNPs (r^2^<0.1) at a harmonic mean FDR-corrected p-value threshold of 0.05 (see Methods), we found the number of genetic loci shared between psychiatric disorders and aggregate SA (N=50) to be approximately twice as high as the number shared with aggregate CT (N=26). SNP rs6224 showed the strongest association between SA and a disorder (SCZ, HMP=3.66×10^−8^), while rs13107325 showing the strongest signal between CT and a disorder (also SCZ, HMP=9.91×10^−13^) (Figure 1, Supplementary Figures 15 and 16 and Supplementary Tables 1 and 2). Seven lead SNPs associated with psychiatric disorders were jointly associated with CT and SA, including rs147317628 for ASD (HMP for CT=1.73×10^−3^, HMP for SA=6.98×10^−4^); rs10433615 for BD (HMP for CT=0.03, HMP for SA=8.44×10^−4^); rs370739771 for PTSD (HMP for CT=0.04, HMP for SA=0.02); and rs60135207 (HMP for CT=0.004, HMP for SA=9.04×10^−4^), rs56219078 (HMP for CT=1.51×10^−4^, HMP for SA=1.38×10^−6^), rs7225002 (HMP for CT=2.18×10^−5^, HMP for SA=1.11×10^−5^) and rs55938136 (HMP for CT=3.28×10^−4^, HMP for SA=7.08×10^−5^) for SCZ (Figure 1, Supplementary Figures 15 and 16 and Supplementary Tables 1 and 2).

Subsequently, we identified specific associations at the brain regional level that contributed to the shared genetic loci identified in the aggregate analysis. Remarkably, at a conjunctional FDR threshold of 0.05, we found concordant and discordant directions of allelic effect between regional brain measures and liability for psychiatric disorders. In this context, a concordant direction of effect implies that greater CT/SA was associated with higher liability. In contrast, a discordant direction of effect implies that greater CT/SA was associated with lower liability. For example, rs10433615 showed a concordant direction of effect in regional CT and SA (Figure 1, Supplementary Figures 15 and 16 and Supplementary Tables 3 and 4), such that a higher genetic vulnerability for BD was associated with greater CT of the medial supplementary motor cortex and greater SA of the insula, lateral temporal and medial prefrontal cortex. In contrast, rs147317628 (associated with ASD), rs370739771 (associated with PTSD), rs60135207 (associated with SCZ) and rs56219078 (associated with SCZ) each showed a concordant direction of effect for CT and a discordant direction of effect for SA, such that greater psychiatric vulnerability was associated with greater CT and lower SA (Figure 1, Supplementary Figures 15 and 16 and Supplementary Tables 3 and 4). However, rs7225002 and rs55938136 (associated with SCZ) showed a discordant direction of effect with CT and concordant direction of effect with SA (Figure 1, Supplementary Figures 15 and 16 and Supplementary Tables 3 and 4).

### Hierarchical gradient of regional brain structures influenced by genetic loci shared with psychiatric disorders

To further delineate the neuroanatomical profile of shared genetic influences between regional brain structure and risk for psychiatric disorders, we used FUMA^50^ to enumerate independent lead SNPs at a conjunctional FDR threshold <0.05 (see Methods). The number of lead SNPs identified across disorders at the regional level was 713 for ADHD (140 for CT, 573 for SA), 126 for ANX (42 for CT, 84 for SA), 593 for ASD (210 for CT, 383 for SA), 754 for BD (333 for CT, 421 for SA), 309 for MDD (115 for CT, 194 for SA), 577 for PTSD (167 for CT, 410 for SA), 8036 for SCZ (2597 for CT, 5439 for SA; Supplementary Tables 5-11). There was substantial heterogeneity across psychiatric disorders in the anatomical pattern of maps of genetic overlap with regional CT and SA (Figure 2A and Supplementary Table 12). Notably, the regions exhibiting a higher number of shared genetic loci with psychiatric disorders included multimodal areas such as the dorsolateral prefrontal cortex, temporo-parietal cortex, and posterior cingulate cortex, as well as unimodal areas such as primary visual and sensorimotor cortex (Figure 2A).

**Figure 2.**
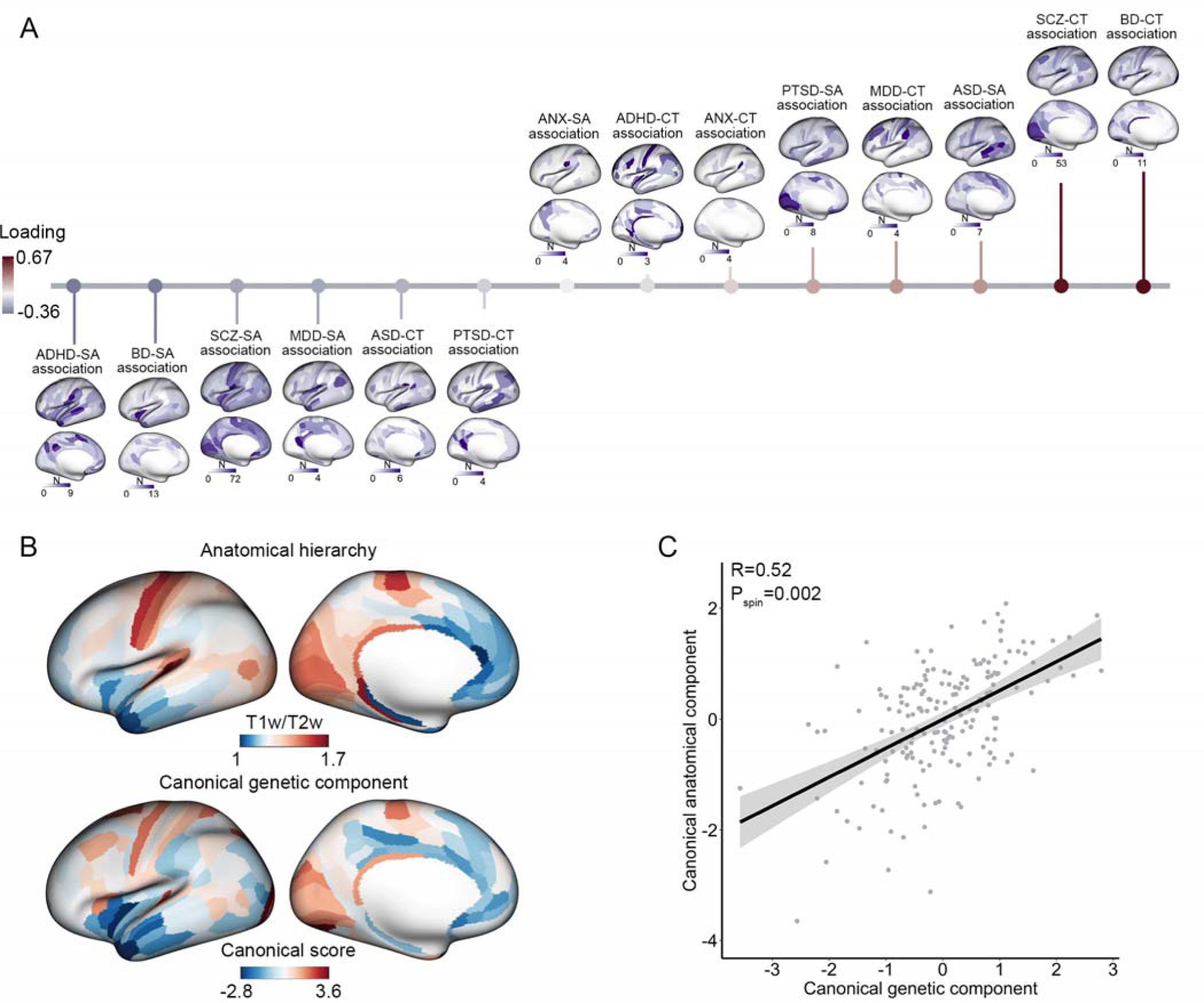
Genetic overlap of psychiatric disorders with the regional brain structure and its association with anatomical hierarchy. (A) The genetic overlap of psychiatric disorders with regional cortical thickness and surface area follows a pattern along an axis of anatomical hierarchy. Each brain map demonstrates the genetic overlap between a specific disorder and regional brain measures. The horizontal color bars indicate the number of shared genetic loci between psychiatric disorders and regional brain structure. These genetic overlap maps exhibit a significant multivariate association with anatomical hierarchy, arranged in ascending order based on their loading values. The length of the vertical lines represents the strength of the loading of each genetic overlap map from the canonical correlation analysis with the map of anatomical hierarchy. A red line indicates positive loadings (a positive contribution to the multivariate association), while a blue bar represents negative loadings (a negative contribution to the multivariate association). (B) The top cortical map illustrates the spatial distribution of T1w/T2w signal intensity, which is inversely related to anatomical hierarchy (i.e., warmer colors correspond to lower hierarchical levels). The bottom cortical map illustrates the canonical component of genetic overlap maps, capturing the joint association between psychiatric disorders and regional cortical structure along a unidimensional spatial component. (C) Scatterplot of the positive correlation between the canonical genetic component and the canonical anatomical component derived from the canonical correlation model. The significance of multivariate association is computed by randomly spatially rotating hierarchical ranks of brain anatomy in 10,000 spin tests.

We sought to determine whether the anatomical pattern of psychiatric-genetic overlap was organized along established anatomical, functional, or evolutionary hierarchies considered to characterize cortical organization^51–54^. For instance, unimodal areas tend to display higher neuron density and greater proportion of feedforward connections originating from the supragranular layer, while multimodal cortices subserving higher-order cognitive processes tend to possess lower neuron density and a predominance of feedback connections originating from the infragranular layer^52,53,55,56^. This neuroanatomical gradient has been quantified by the ratio of T1-weighted to T2-weighted MRI images (T1w/T2w), which is negatively correlated with anatomical hierarchy^51^. We used a multivariate approach, canonical correlation analysis (CCA), to interrogate the association between this anatomical gradient and the anatomical pattern of psychiatry-genetic overlap, defined by the number of shared independent lead SNPs in each of the conjunctional FDR analyses. Remarkably, there was a significant association between canonical anatomical and psychiatric-genetic components derived from CCA (r=0.52, p_spin_=0.002, Figure 2). In contrast, there was no statistically significant alignment with canonical maps of functional organizational derived from functional MRI studies^57^ (p_spin_=0.67, Supplementary Figure 17) or a map of the evolutionary expansion of the cortex^58^ (p_spin_=0.41, Supplementary Figure 17), suggesting a relatively specific alignment of the cortical pattern of psychiatric-genetic overlap with the microstructural, anatomical hierarchy quantified by T1w/T2w. By further examining the loadings of the canonical psychiatric-genetic component from the CCA with T1w/T2w (see Methods), we further observed a gradient of shared genetic variants between psychiatric disorders and cortical anatomy. For instance, genetic overlap between CT and SCZ/BD was observed predominantly in lower-order cortex, while shared genetic overlap SA and SCZ/BD was observed in higher-order cortex (Figure 2 and Supplementary Table 13). This alignment suggests that the variance in genetic overlap of psychiatric disorders with brain structures can be explained, at least in part, by the underlying microstructural, anatomical hierarchy.

### Mixture of concordant and discordant genetic associations with regional brain structure

By harmonizing the direction of effect for all risk alleles, we directly assessed whether the risk allele of each lead SNPs in the conjunctional FDR analysis had a concordant (increased CT or SA) or discordant (decreased CT or SA) direction of effect on regional brain structure. Homogeneity in the directions of effect across shared genetic loci would support common pathophysiological underpinnings between the brain structure and psychiatric disorders, whereas heterogeneity, particularly in the directions of effect of SNPs affecting the same brain region, would support distinct pathophysiological mechanisms (Supplementary Tables 5-11). Cumulatively, by mapping the differences in the number of genetic loci displaying concordant and discordant directions of effect within each brain measure, our results indicate a mixture of more homogenous and more heterogeneous directions of effect across psychiatric disorders (Figure 3 and Supplementary Table 14).

**Figure 3.**
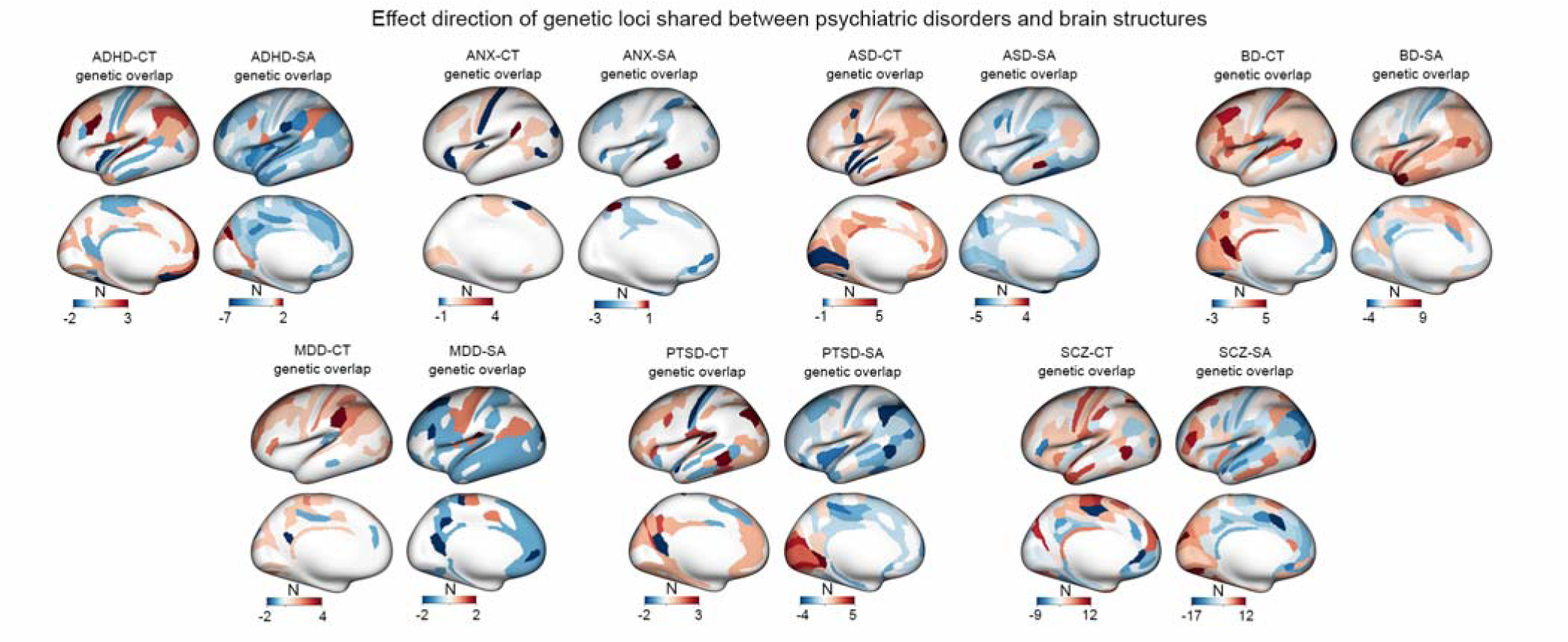
Effect direction of shared loci between psychiatric disorders and brain structure. The cortical maps illustrate the disparity between the number of genetic loci exhibiting a concordant direction of effect and those displaying a discordant direction of effect between psychiatric disorders and brain structure. In the maps, red indicates brain regions where the number of genetic loci with a concordant direction of effect (i.e., the risk allele is associated with greater surface area or cortical thickness) exceeds those with an discordant direction of effect (i.e., the risk allele is associated with lower surface area or cortical thickness), while blue represents brain regions where the number of genetic loci with a discordant direction of effect is higher than those with a concordant direction of effect.

For ADHD, ANX, ASD, MDD and PTSD, results were skewed towards concordant directions of effect for CT and discordant directions of effect for SA (Figure 3 and Supplementary Table 14). Specifically, in the case of ADHD, shared genetic loci with regional CT measures were 75% concordant, while shared genetic loci with regional SA measures were 91% discordant. For example, 3 ADHD lead SNPs were associated with greater thickness of the dorsolateral prefrontal cortex (rs945501, rs76338508 and rs6789751), and 7 ADHD lead SNPs were associated with lower SA of the parietal operculum (rs3791102, rs4367080, rs223498, rs12054920, rs10956838, rs7014426, rs1585897, rs77653640, rs9931589). In ANX, shared genetic loci were 70% concordant for CT and 94% discordant for SA. For example, 4 lead SNPs were associated with greater thickness of perisylvian cortex (rs631653, rs13263075, rs11112594, and rs79691782), and 3 lead SNPs were associated with lower SA of the superior parietal cortex (rs34402524, rs4613794 and rs13263075). In ASD, shared genetic loci were 91% concordant for CT and 92% discordant for SA. For example, 5 lead SNPs were associated with greater thickness of the inferior temporal cortex (rs6543224, rs56842404, rs2079227, rs147317628, rs56305452), and 5 lead SNPs were associated with lower SA of the perirhinal cortex (rs10188273, rs11147235, rs147317628, rs56305452 and rs35570734). In MDD, shared genetic loci were 87% concordant for CT and 90% discordant for SA. For example, 4 lead SNPs were associated with greater thickness of the supramarginal cortex (rs4437406, rs943840, rs10142766 and rs911552). In PTSD, shared genetic loci were 83% concordant for CT and 80% discordant for SA. For example, 3 lead SNPS were associated with greater thickness in the inferior temporal cortex (rs7524494, rs139677376 and rs370739771), and 4 lead SNPs were associated with lower SA in the inferior parietal cortex (rs112416932, rs9375138, rs2049114, rs370739771).

Different patterns in the direction of effect were observed for BD and SCZ. In BD, shared genetic loci were 85% concordant for CT but also 78% concordant for SA. Most notably, for 9 of lead SNPS jointly associated BPD, the risk allele was also associated with greater SA of the temporal pole (rs2258734, rs2083180, rs6997571, rs12351417, rs4930184, rs10412176, rs6088607, rs2143943 and rs5753507). In contrast, the directions of effect in SCZ were relatively evenly distributed -- shared genetic loci were 57% discordant for CT and 62% concordant for SA in their direction of effect (Figure 3 and Supplementary Table 14). These results are consistent with the heterogeneity of findings in decades of research on regional alterations in brain structure in SCZ, which has previously been attributed to the proposed existence of multiple neurobiological or genetic subtypes of SCZ^59^.

### Significant pleiotropy obscured by absent genetic correlations between brain structure and psychiatric disorders

Heterogeneous directions of effect between a psychiatric disorder and a neuroimaging phenotype may result in pleiotropic effects being underestimated by the genetic correlation between two traits. To test this hypothesis, we used LDSC to estimate the genetic correlations between each pair of psychiatric disorders and measures of CT and SA using GWAS summary statistics. Among SA measures, 76 regions exhibited significant negative genetic correlations with brain disorders after correction for FDR at p<0.05 (Supplementary Figure 18, Supplementary Tables 15 and 16). The large majority of these correlations were found with ADHD (negative genetic correlations with 69 regional measures of SA across parietal, temporal, dorsolateral prefrontal, and medial prefrontal cortex). In addition, MDD was negatively genetically correlated with SA in 5 regions in the inferior temporal cortex, and PTSD was negatively correlated with 2 regions in the superior parietal cortex. For the 76 regional SA measures with statistically significant genetic correlations with ADHD, MDD, or PTSD, conjunctional FDR analysis identified a total of 255 lead SNPs (246 associated with ADHD, 5 lead SNPs associated with MDD, and 4 lead SNPs associated with PTSD; Supplementary Table 19). Consistent with the findings of the global genetic correlation analysis, a significant proportion of these lead SNPs (204 out of 255) exhibited a discordant direction of effect such that the risk allele was associated with lower SA.

None of the psychiatric disorders showed a significant global genetic correlation with CT, nor did any of SCZ, BPD, ASD, or ANX show statistically significant genetic correlations with SA (Supplementary Figure 19 and Supplementary Tables 17 and 18). Among the regional phenotypes that did not show significant genetic correlations with any psychiatric disorder (104 regional SA measures and 180 regional CT measures), the conjunctional FDR analysis identified a total of 10,853 lead variants that were shared across different psychiatric disorders. Remarkably, 50.51% (5,482 out of 10,853) of these shared genetic loci exhibited a concordant direction of effect, while 49.49% exhibited a discordant direction of effect. These results confirm that while significant genetic correlations between imaging phenotypes and psychiatric disorders are indicative of pleiotropy, significant pleiotropy exists even in the absence of genetic correlations. Analytic approaches such as conjunctional FDR provide insights into the bi-directional relationships obscured by conventional genetic correlation analyses.

### A subset of genetic loci that influence brain structure and multiple psychiatric disorders

Given significant evidence of pleiotropy between psychiatric disorders and brain structure, and prior evidence of genetic correlations among psychiatric disorders, we sought to identify specific genetic loci associated with regional CT or SA as well as multiple psychiatric disorders. Overlapping patterns of pleiotropy with brain structure would support a potential common underlying biological mechanism influencing CT/SA among disorders. In support of this hypothesis, our analysis revealed a total of 160 lead SNPs that were shared among psychiatric disorders. Out of these shared SNPs, 132 exhibited either partial or complete overlap in terms of their CT or SA profiles between two disorders (Figure 4 and Supplementary Figures 20-30).

**Figure 4.**
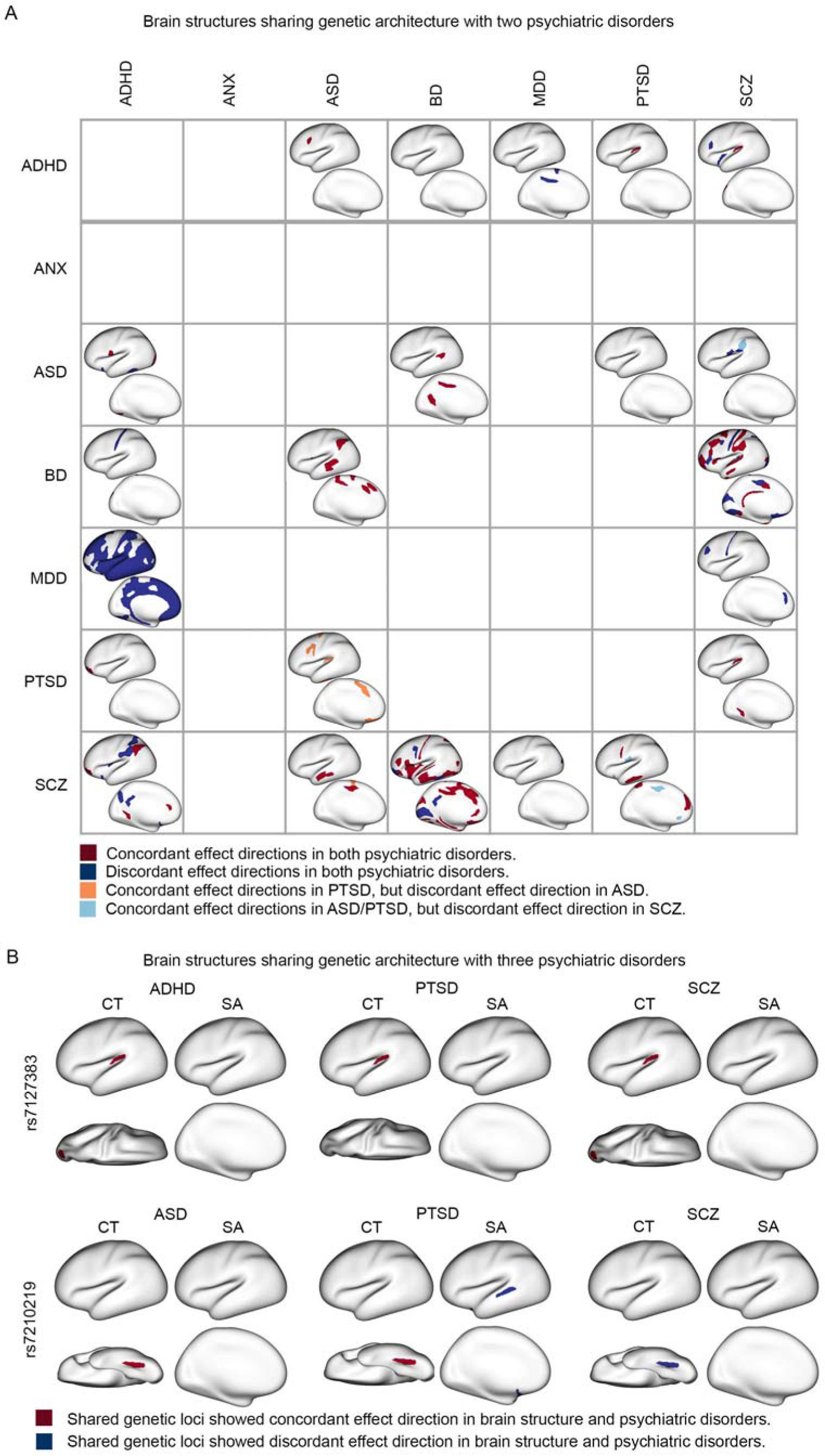
Overlapping genetic effect of psychiatric disorders on brain structures. (A) The impact of shared genetic loci on brain structure for two psychiatric disorders is depicted. The lower left triangle illustrates the overlapping SA measures between the shared genetic loci for the two disorders. The upper right triangle illustrates the overlapping CT measures between the shared genetic loci for the two disorders. Dark red represents brain regions where the effect directions of shared genetic loci are consistent in both disorders. Dark blue represents brain regions where the effect directions of shared genetic loci are opposite in the two disorders. Orange and light blue represent brain regions where the effect directions of shared genetic loci are concordant in one disorder but concordant in the other disorder. (B) The overlapping brain patterns between shared genetic loci for three psychiatric disorders are shown. Red represents brain regions where the effect direction of genetic association is consistent in both the brain region and the respective psychiatric disorder. Blue represents brain regions where the effect direction of genetic association is negative in both the brain region and the psychiatric disorder.

Out of the 132 SNPs with overlapping anatomical profiles in the conjunctional FDR analysis with two disorders, 113 SNPs were consistent in terms of their direction of effect for both disorders. For example, the risk allele of rs3851274 for both ADHD and ASD was associated with greater thickness of the dorsolateral prefrontal cortex (Figure 4 and Supplementary Figure 20). The risk allele of rs1585897 for both ADHD and MDD was associated with lower SA in multiple areas of association cortex in (Figure 4 and Supplementary Figure 22). The risk alleles of 6 lead SNPs common to both ASD and BD (rs9401593, rs1906252, rs9375188, rs17814604, rs1487441, and rs1487445) were associated with greater CT and SA in superior temporal, supramarginal, anterior cingulate, and motor cortex (Figure 4 and Supplementary Figure 25). Consistent with their high genetic correlation, BD and SCZ showed the most anatomic overlap in genetic loci with pleiotropic effects on brain structure (69 lead SNPs in total). For example, the risk alleles of 9 SNPs common to both BD and SCZ (rs2169895, rs3769484, rs514413, rs55765017, rs55931635, rs6461049, rs6984358, rs871925, rs295140) were associated with lower SA and CT of the primary cortex, including somatosensory and visual cortex (Figure 4 and supplementary Figure 28). Finally, the risk allele of one lead SNP -- rs7127383, common to ADHD, PTSD, and SCZ -- was implicated in three disorders with a consistent direction of effect. The risk allele was associated with greater thickness of the auditory cortex in the conjunctional FDR analysis for each of the three disorders (Figure 4) and with greater thickness in the lateral visual cortex in the conjunctional FDR analysis of ADHD and SCZ (Figure 4).

In contrast, 19 SNPs with overlapping anatomical profiles with two disorders showed divergent directions of effect. For instance, for 9 risk loci common to ASD and PTSD (rs2445956, rs11123965, rs264975, rs2198234, rs1673468, rs264963, rs10188273, rs2084814, rs1869073), the risk alleles were associated with lower SA for the somatosensory, anterior cingulate and orbitofrontal cortex in the conjunctional FDR analysis with ASD (Figure 4 and Supplementary Figure 26). However, the risk alleles of these SNPs were associated with higher SA of the same cortical areas in the conjunctional FDR analysis with PTSD (Figure 4 and Supplementary Figure 26). In addition, one lead SNP -- rs7210219, common to ASD, PTSD, and SCZ – was implicated in three disorders with a mixed direction of effect. The risk allele was associated with greater thickness of the fusiform gyrus in the conjunctional FDR analysis of both ASD and PTSD but lower thickness of this gyrus in the conjunctional FDR analysis of SCZ (Figure 4).

As noted above, two lead SNPs were implicated in the conjunctional FDR analysis with brain structure for three distinct disorders (rs7127383 and rs7210219). At the conjunctional FDR threshold of 0.05, no genetic loci were implicated in four or more distinct disorders. Moreover, we did not find any common genetic loci between ANX and the other psychiatric disorders in the conjunctional FDR analyses with brain structures (Figure 4).

### Functional annotation of shared genetic loci between psychiatric disorders and brain structures

We conducted functional annotation for each lead genetic variant that was shared between brain disorders and brain structures in the conjunctional FDR analysis. The highest proportion of exonic variants was observed in SCZ, accounting for 3.1% of the shared variants (Figure 5A and Supplementary Tables 20 and 17). *CUL9* contains three exonic variants jointly associated with SCZ and CT (rs6917902, rs2273709, rs9472022). The latter two SNPs were both associated with greater thickness of fusiform cortex (Supplementary Table 11). The majority of shared genetic variants, across all psychiatric disorders, were primarily located in intronic or intergenic regions that likely play a role in modulating gene expression and chromatin accessibility (Figure 5A and Supplementary Tables 20-27). Quantified by the proportion of genetic loci identified in the disorder-brain conjunctional FDR analysis containing expression quantitative trait loci (eQTLs) or chromatin accessibility QTL (caQTLs) identified with FUMA^50^ (Supplementary Tables 21-27), there was a particularly high degree of enrichment for regulatory genomic regions in BD (79%), ASD (71%), ANX (71%), and SCZ (67%),

**Figure 5.**
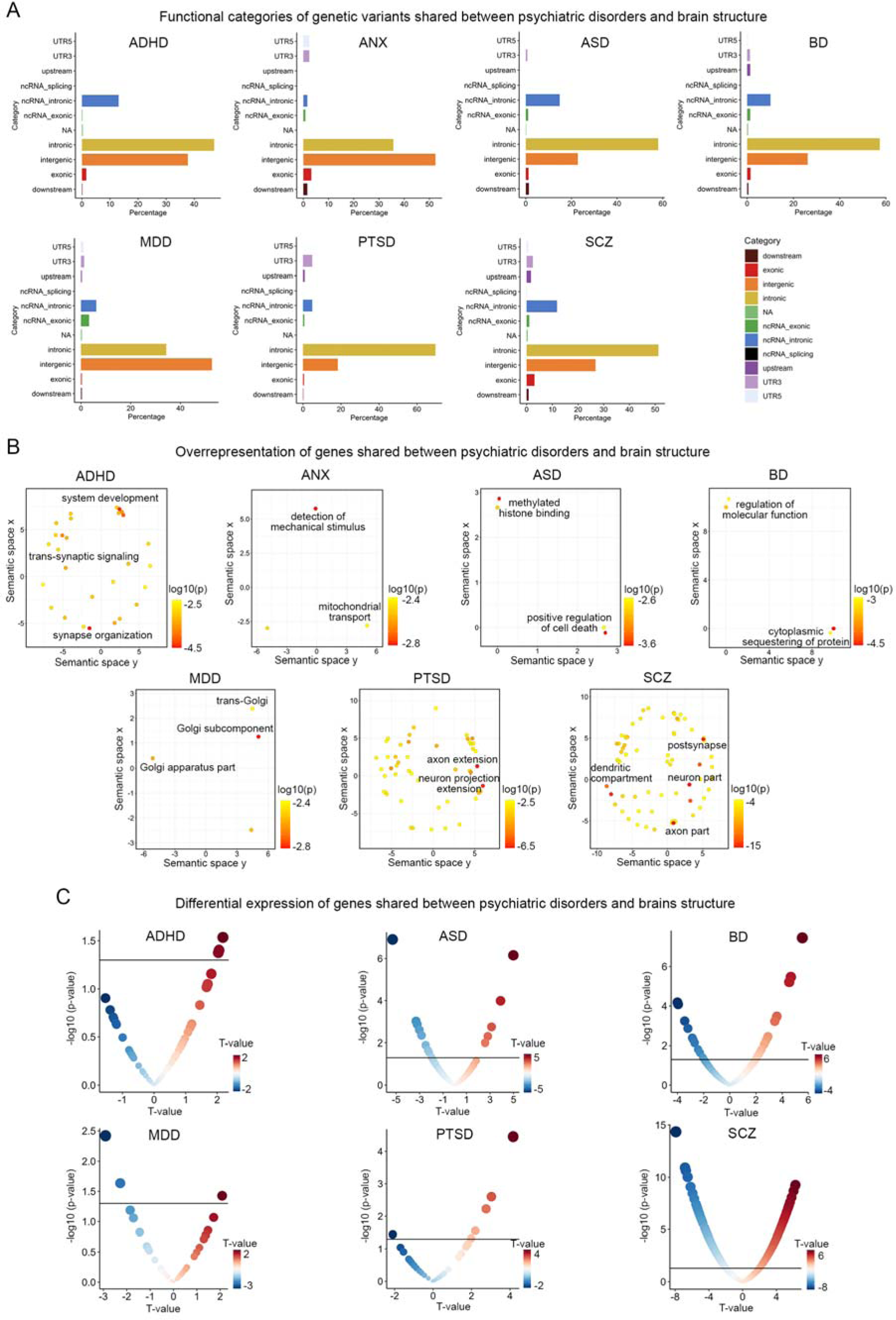
Functional annotation and differential expression of shared genes between psychiatric disorders and brain structures. (A) Functional categories of lead genetic variants shared between psychiatric disorders and brain structures identified through conjunctional FDR analysis. The x-axis represents the proportion of shared genetic variants within each specific functional category, while the y-axis represents different functional categories. Each color corresponds to a particular functional category. (B) Overrepresentation analysis of genes located nearest to the shared genetic loci between psychiatric disorders and brain structures, presenting the results in a two-dimensional space using REVIGO. REVIGO reduces the dimensionality of gene-ontology terms based on pairwise semantic similarities. Red indicates biological terms with a lower p-value in the overrepresentation test. (C) Differentially expressed genes that overlap with genetic loci identified in the conjunctional FDR analysis of GWAS of psychiatric disorders and cortical brain structure. The horizontal black lines represent the significance threshold of p<0.05. Red dots indicate genes with up-regulated expression in individuals with the disorder compared to controls, while blue dots indicate genes with down-regulated expression. The size of the dots is scaled according to the absolute value of the *t*-values.

Next, we mapped each shared independent lead SNP to the closest gene based its genomic position (see Methods). To identify pathways and biological processes associated with the shared gene between psychiatric disorders and CT and SA, we performed overrepresentation analyses of gene ontology terms using ConsensusPathDB^60^ (see methods). A total of 594 molecular processes exhibited significant overrepresentation (FDR p<0.05) across all brain disorders (Figure 5B and Supplementary Table 28). The most significant processes implicated in the genetic association of ADHD and brain structures was “synaptic organization” (hypergeometric test p=1.87×10^−5^). For example, the three most significant lead SNPs were in *ST3GAL3* (rs35891206, rs2527776 and rs803678), which is considered to mediate synaptic plasticity and cognitive deficits in knockout mouse models^61^. Genes jointly associated with brain structures and ASD showed the strongest enrichment in “methylated histone binding” (p=0.001). rs147317628, which is jointly associated with ASD and 62% of brain measures, is an intronic variant in *KANSL1*, a crucial chromatin modifier implicated in the pathogenesis of ASD^62^. The top-ranked gene set involving genes jointly associated with brain structure and BD is “cytoplasmic sequestering of protein” (p=2.46×10^−5^), which is important for the maintenance of protein transport and protein location within mammalian cells. For example, ANKRD23, which is extensively implicated in the genetic overlap between BD and CT, is a scaffold protein to regulate the assembly of voltage-gated sodium channels^63,64^. Top molecular processes influencing the genetic association between MDD and brain structure were predominantly involved in Golgi complex cellular components including “trans-Gogli network” (p=0.005). Processes identified for SCZ encompassed fundamental components of neural architecture including “post-synapse” (p=9.82×10^−16^), “dendrite” (p=1.46×10^−13^), and “axon” (p=5.26×10^−8^). For example, the strongest association between SCZ and SA is observed with the intronic variant rs6224 of *FUSIN*, a gene prominently expressed in the brain and exhibiting disrupted expression in psychotic brains^65^. This protein is also known to mediate a range of neural processes, including synaptic plasticity, neural proliferation, dendrite development, neural survival and death^65^. Notably, for the association between PTSD and brain structure, we found novel evidence of biological processes related to different aspects of neuronal projection including “axon extension” (p=2.51×10^−7^) and “neuron projection extension” (p=2.29×10^−6^). Finally, no significant pathways were identified for ANX at FDR threshold<0.05, but at a more lenient 0.1 threshold, we observed an enrichment for mitochondrial components including “mitochondrial transport” (p=0.004).

Utilizing publicly available transcriptomic profiles derived from postmortem brain case-control studies^66–68^ (see Methods), we next determined if there was overlap between the genes closest to shared independent lead SNPs identified through the conjunctional FDR analysis and differentially expressed genes (DEGs) in the corresponding brain disorder. This overlap was higher in ASD (16 DEGs), BD (21 DEGs), and SCZ (300 DEGs) – in line with larger sample sizes in case-control studies of these disorders as well as higher degree of enrichment for regulatory genomic regions noted above – compared to ADHD (3 DEGs), MDD (3 DEGs), or PTSD (6 DEGs) (differential expression data was unavailable for ANX) (Figure 5C and Supplementary Tables 29-34). For example, a lead SNP jointly associated with ASD risk and with lower CT in lateral prefrontal cortex (rs2084200; conjunctional FDR=0.04; Supplementary Table 7) is an intronic SNP and an eQTL of *WDPCP*, a gene involved in cell polarity that exhibited lower expression level in individuals with ASD compared to controls (p=0.001, t=-3.31). The gene *LINGO1* showed pronounced lower expression in postmortem cortex of individuals with SCZ (p=8.19×10^−10^, t=-6.18), and SCZ-brain FDR analysis showed two lead SNPs (rs28661259, rs883669) intronic to *LINGO1* that were independently associated with greater CT of the cingulate cortex (FDR=0.03) and greater SA of the orbitofrontal cortex (FDR=0.048; Supplementary Table 11). LINGO1 was also a core gene of the overrepresented pathway involving neural differentiation identified in functional annotation of SCZ-brain results (Figure 5b). Moreover, a lead SNP jointly associated with BD risk and increased CT of the dorsolateral prefrontal cortex (rs72724966; conjunctional FDR=0.04, Supplementary Table 8) was also proximal to *LINGO2,* a gene showing pronounced increased expression in postmortem cortex of individuals with BD (p=3.38×10^−6^, t=4.66; Supplementary Table 34).

## Discussion

The present study expands and clarifies the landscape of genetic relationships between psychiatric disorders and the anatomy of the human cerebral cortex, using publicly available data from approximately 700,000 participants. We employed conjunctional FDR analysis to demonstrate extensive pleiotropy between seven psychiatric disorders and MRI-based quantifications of regional CT and SA, patterned along an axis of anatomical hierarchy that contained poles in primary and association cortices. Notably, psychiatric risk alleles exhibited diverse directions of effect on CT and SA, helping to explain relatively low brain-disorder global genetic correlations. Via functional annotation of genetic loci with shared influences on brain structure and psychiatric liability, we uncovered involvement in neurobiological, metabolic, and transcriptomic regulation pathways, including genes that are differentially expressed in post-mortem cortex of individuals with psychiatric disorders. Collectively, our findings provide valuable insights into the underlying neuropathological pathways that connect genetic risk, the anatomical organization of the cerebral cortex, and psychopathology.

Differences between individuals in cortical anatomy have been observed for decades to be associated with differences in cognitive function, mental health, and various dimensions of psychopathology^69–71^. There have been many attempts to establish brain MRI measures as endophenotypes for psychiatric disorders, and previous studies have primarily investigated phenotypic associations in patients and unaffected relatives^72,73^. Here, we used a novel Bayesian-based approach to identify genetic variants that are jointly associated with both psychiatric disorders and brain structural variations. Compared to previous studies that have aggregated genetic effects associated with specific psychiatric disorders into a single polygenic score^34^, our approach provides comprehensive and systematic evidence for the shared genetic mechanisms by examining the specific genetic variants and their directions of effect. Our findings indicate that previously observed alterations in brain morphology in individuals with psychiatric disorders may be influenced, in part, by shared genetic determinants. One area for future research is to develop pleiotropy-enriched polygenic scores that incorporate the prioritized genetic loci identified in the conjunctional FDR analysis.

Our multivariate analysis revealed that genetic loci that increase risk for psychopathology also influence cortical anatomy along an axis of anatomical hierarchy. For example, we observed that CT’s genetic overlap with liability for ADHD, BD, and SCZ was aligned with regions of the primary cortex such as visual cortex and sensorimotor cortex. On the other hand, SA’s genetic overlap with ADHD, BD, and MDD was aligned with regions of association cortex including insula, anterior temporal cortex, and medial prefrontal cortex. This hierarchical organization of genetic influence on brain structure is supported by neuroanatomical evidence across different species. Regions lower in the hierarchy, such as the primary cortex, are characterized by specialized physiological and functional properties, including a high density of neurons and a significant proportion of supragranular-originating neurons that contribute to feedforward information processing for externally oriented perception^52,53,55^. In contrast, regions higher in the hierarchy, such as the association cortex, exhibit less differentiation in cytoarchitecture, including lower neuronal density and a predominance of infragranular-originating connectivity involved in feedback communication for internally focused cognitive functions^56^. Moreover, neuroimaging-transcriptomics associations have demonstrated that the hierarchical organization of the human brain’s anatomy is shaped by gradients of gene expression, which are related to cell type specificity and laminar enrichment^51^. Building upon this evidence, we hypothesize that risk for psychiatric disorders is influenced by the transcriptional regulation of genes within specific cell types and cortical layers that also influence cortical brain structure. For example, we identified four SNPs (rs10261780, rs66571810, rs17137124, and rs10261780) that jointly associate with ADHD and the thickness of the sensorimotor cortex in our conjunctional FDR analysis. These intronic variants are located within the *FOXP2* gene, which is predominantly expressed in the inner cortical layers^74–76^ and plays a regulatory role in motor circuits^77,78^. We speculate that these ADHD risk variants may modulate the laminar expression of *FOXP2*, thereby influencing the cytoarchitecture of somatomotor cortices.

We found that risk alleles for psychiatric disorders exhibited mixed directions of effect on cortical anatomy. For the majority of disorders, risk alleles tended to be associated with higher CT or with lower SA. These pleiotropic genetic loci may exert their influence on both the development of brain disorders and cortical morphology through common intermediate endophenotypes, such as regulation of cellular gene expression and molecular signal transmission. A distinctive pattern was observed for SCZ, such that risk alleles showed a balanced mixture of associations with higher and lower CT or higher and lower SA. This finding is noteworthy considering the inconsistent and equivocal results reported in previous studies investigating alterations in regional morphology in individuals with SCZ^14,79–81^. Our results shed light on the underlying mechanisms of the heterogeneous neuroanatomical changes associated with SCZ, highlighting the role of genetic pleiotropy. It is important to acknowledge that characterizing pleiotropic loci between a specific psychiatric disorder and cortical structure is a complex task due to the influence of external environmental variables and factors beyond disease-specific genetics. Clinical heterogeneity and comorbidity, for instance, pose significant challenges in this context. These factors can contribute to the variability in neuroanatomical changes observed in disorders such as SCZ. To address these challenges and gain further insights, future investigations should consider defining subgroups within the pleiotropic loci based on concordant and discordant directions of effect, to identify common molecular signatures of specific morphological alterations.

The mixed directions of effect for pleiotropic loci point to limitations on the utility of global genetic correlation as a measure of genetic overlap between neuroimaging measures and psychiatric disorders. In the LDSC-based genetic correlation analysis, we identified a negative correlation between three psychiatric disorders and the SA of the association cortex. Within these three disorder-brain pairs, our conjunctional FDR analysis identified a total of 255 independent lead variants, of which 80% exhibited a discordant direction of effect (i.e., the risk allele was associated with lower SA). These findings are consistent with LDSC, as an approach to assess global genetic correlation, being sensitive to correlations across SNPs with a consistent direction of effect^29^. While this method provides valuable insights into the genetic overlap between phenotypes, it does not capture the full extent of genetic interactions, as genetic variants with discordant and concordant directions of effect can “cancel out”. By leveraging pleiotropy enrichment and conditional re-ranking statistics to enhance statistical power, conjunctional FDR identified shared genetic variants with effects in both directions across all disorder-brain pairs^37^. It is worth noting that this approach also mitigates bias towards the most highly powered GWAS, ensuring that the identified genetic variants were not disproportionately influenced by any particular trait^36^.

For most genetic risk loci that were shared across multiple psychiatric disorders, our results demonstrated convergent associations with cortical morphology. For example, 83% of shared genetic loci exhibited partially or completely overlapping brain maps of statistical associations with cortical morphology, and 86% showed consistent directions of effect on the same brain measures. Strikingly however, a subset of shared genetic loci displayed antagonistic effect directions on the same morphological measures across disorders. For example, genetic variants associated with ASD and PTSD exhibited opposite effect directions on the SA of the dorsal prefrontal cortex and anterior cingulate cortex. While the relationship between ASD and PTSD in terms of brain structural vulnerability is still relatively unexplored^82^, previous large-scale neuroimaging consortium studies have demonstrated opposite patterns of association between these two disorders and the dorsal prefrontal cortex and anterior cingulate cortex^13,83^. This supports the hypothesis that shared genetic loci between ASD and PTSD contribute to neuropathological changes in these regions through divergent effect directions. By contrast, the risk allelic effect of rs7210219 for both ASD and PTSD had a concordant association with the thickness of the fusiform cortex, but a discordant association was observed for SCZ. Collectively, our findings support substantially overlapping neurogenetic mechanisms across mental disorders but also suggest the existence of a smaller subset of distinct neurogenetic pathways that are not shared across disorders. It is worth noting that the total number of shared loci identified (and thus power to detect convergent or divergence associations with cortical morphology) is influenced both by the heritability of each trait and by the sample size of existing GWAS. For instance, the relatively low sample size of existing GWAS for ANX likely explains the lack of shared genetic variants observed between ANX and other psychiatric disorders.

Functional annotation of overlapping genetic loci likewise revealed evidence for a mixture of common and distinct pathways among psychiatric disorders that also influence neuroimaging measures. Unsurprisingly, common pathways implicated processes related to synaptic organization, neuronal differentiation and growth, and cellular metabolism. Consistent with previous GWAS studies for complex traits^2,21,22,84^, our results also highlighted the significance of intronic and intergenic loci that function in chromatin regulation and the regulation of gene expression in the human brain.^85^ ^86–88^. Risk loci for ASD, BD, and SCZ had relatively high proportion of overlap with eQTL/chromatin interaction loci and with genes that were differentially expressed in post-mortem patient brain tissue. On the other hand, risk loci for ADHD, MDD, and PTSD displayed relatively weaker overlap with differentially expressed genes. It should be noted that some discrepancies may arise from the incomplete characterization of disease transcriptomic profiles, as transcriptomic data were largely limited to the dorsolateral prefrontal cortex and anterior cingulate cortex, though recent evidence supports relatively consistent differential expression patterns across most cortical areas^89^. As the availability of transcriptomic data from patient tissue continues to expand, there is an unprecedented opportunity for future work to systematically explore the molecular regulatory mechanisms of shared genetic loci between psychopathology and brain structure.

### Limitations

Several limitations should be acknowledged in the present study. Firstly, although our conjunctional FDR analysis effectively captured pleiotropic loci between psychopathology and cortical structure, assessing causality remains challenging for genetic variants exhibiting small effect sizes on complex traits. Therefore, it remains uncertain whether the development of psychiatric disorders is influenced by the modification of shared genetic loci affecting brain anatomical organization, or whether alterations in brain structure are caused by genetic contributions to these disorders. Future investigations utilizing longitudinal study designs will be necessary to elucidate such causal relationships. Secondly, psychiatric disorders frequently exhibit high comorbidity^90,91^, with approximately 40% of individuals meeting criteria for one diagnosis also meeting criteria for at least one additional disorder^92^. Our study does not rule out the possibility that GWAS cohorts may include individuals experiencing multiple psychiatric symptoms or undergoing diagnostic transitions, such as individuals with MDD transitioning to other diagnostic categories like BD after their inclusion in the study. Thirdly, our analysis predominantly focused on identifying common genetic architecture based on GWAS of individuals with European ancestry, limiting the generalizability of our findings to populations with different genetic ancestries and ethnic backgrounds. Future research should aim to broaden the scope of investigation to include diverse populations to enhance the external validity and applicability of our results. Finally, we are unaware of additional accessible large-scale imaging-genetic and psychiatric GWAS data beyond the data we used for our main analyses, which prevents us from directly assessing replicability. While our study contributes valuable insights into the genetic overlap between psychiatric disorders and brain structures, independent replication using additional datasets is necessary to validate and confirm the observed associations genetics, psychiatric disorders, and brain structure.

## Conclusions

Our study leveraged pleiotropy-informed genetic analysis to delineate previously unreported genetic overlap between 7 psychiatric disorders and 360 neuroimaging measures of cortical morphometry. Psychiatric risk alleles had mixed directions of effect on the thickness and surface area of specific regions of the cerebral cortex, with mostly convergent but in some cases divergent neuroanatomical profiles associated with risk alleles across disorders. Multi-scale functional annotation of this polygenic architecture implicated gene expression regulatory networks, including SNPs that influence genes that are differentially expressed in psychiatric condition. Taken together, these findings represent a significant advance in our understanding of the landscape of shared genetic architecture between psychopathology and cortical brain structure.

## Material and Methods

### GWAS data

For neuroimaging measures, we conducted univariate GWAS analyses for 180 averaged measures of cortical thickness and surface area across hemispheres using data from two cohorts: 31,797 subjects of European ancestry from the UK Biobank and 4,866 subjects of European ancestry from the ABCD study^23^. Cortical regions were defined using the multi-modal Human Connectome Project parcellation^48^, and CT and SA were estimated using FreeSurfer 6.0^93^. To combine the results across cohorts, we performed inverse-variance weighted meta-analyses. The resulting meta-analyzed GWAS summary statistics, comprising a total of 36,663 individuals, were then utilized for subsequent conjunctional FDR analysis. Additional details regarding these GWAS analyses have been described in a separate publication^23^.

For psychiatric disorders, we used summary statistics from previously published GWAS of ADHD (N=53,293)^41^, ANX (N=17,310)^42^, ASD (N=46,350)^43^, BD (N=51,710)^44^, MDD (N=143,265)^45^, PTSD (N=214,408)^46^ and SCZ (N=130,644)^47^. Given that the conditional FDR analysis is sensitive to the overlapping samples, none of the psychiatric GWAS used data from UK Biobank or ABCD cohorts^37^. Due to differences in LD structure across ancestries, all cohorts were limited to individuals of European descent. We did not include obsessive-compulsive disorder in our study because the sample size of the case-control design (Ncase=2,688, Ncontrol=7,037) was deemed insufficient^94^.

### Global genetic correlation

Using bivariate LDSC^28,29^, we estimated the genetic correlation between each pair of brain disorders and regional CT or SA based on GWAS described above. Briefly, GWAS summary statistics were regressed on LD scores derived from the 1000 Genomes European dataset^95^, to test whether the genetic association from the two GWAS datasets differed from what would be expected by chance. To assess the significance of the genetic correlations, we employed FDR^96^ correction for multiple comparisons at a corrected threshold of p<0.05.

### Conjunctional FDR analysis

The pleioFDR toolbox^37^ was used to perform conjunctional FDR analysis, to investigate shared genetic variants between regional CT or SA and brain disorders. This approach enhances the statistical power to detect overlapping genetic loci by leveraging cross-trait SNP enrichment within an empirical Bayesian statistical framework. To mitigate potential inflation of FDR estimates, certain genomic regions were excluded from the analysis due to their complex LD structure. Specifically, SNPs near the extended major histocompatibility complex region (Chr6:25119106-33854733) and chromosome 8p23.1 (Chr8:7200000-12500000) were excluded. For each pair of phenotypes, consisting of a psychiatric disorder and regional measure of CT or SA, the conditional FDR analysis involved re-ranking the test statistics of the brain measure conditioned on its association with the disorder (i.e., *FDR (brain | disorder)*). Conditional FDR values, indicating the significance of association with the brain measure, were recalculated based on this conditioning. Similarly, by reversing the roles of the brain measure and brain disorder, conditional FDR values assessing the significance of SNP associations with the brain disorder were obtained (i.e., *FDR (disorder | brain)*). In contrast to the standard FDR ranking, which provides the same ordering of SNPs, re-ordering SNPs based on the conditional FDR analysis results in a different ranking when the primary and auxiliary phenotypes are genetically related. The conjunctional FDR value, determined as the maximum of the two conditional FDR values in both directions (i.e., *FDR (brain | disorder) and FDR (disorder | brain)*), conservatively estimates the posterior probability that a SNP has no association with either trait, given that its p-values for association with both phenotypes are as small as or smaller than the observed p-values. The significance threshold for shared genetic loci in each disorder-brain pair was set at a conjunctional FDR<0.05. Furthermore, we assessed the directional effects of the shared genetic loci for each disorder-brain pair by comparing their z-scores.

To address the challenge of multiple testing correction, we employed an aggregate analysis strategy, focusing on each psychiatric disorder independently. For each SNP, we calculated the HMP value^49^ by considering the 180 conjunctional FDR values derived from the conjunctional analyses of a specific psychiatric disorder and regional measures of CT or SA. Significance of SNPs was determined based on the HMP threshold of 0.05.

### Definition of independent lead SNPs

To delineate independent genetic loci within the aggregate and separate conjunctional FDR analysis, we utilized FUMA, an online platform designed for functional annotation of GWAS results^50^. Specifically, based on the pre-calculated LD structure from the 1000 Genomes European reference panel^95^, SNPs with genome-wide conjunctional FDR or HMP values <0.05 that had LD r^2^<0.6 with any other SNPs were identified. Independent lead SNPs were also defined among them as having low LD (r^2^<0.1) with any other SNPs. If LD blocks of significant SNPs are located within 250 kb of each other, they were merged into one genomic locus.

### Functional annotations of lead SNPs

We used ANNOVAR^97^ to identify the functional categories of the lead SNPs shared between psychiatric disorders and CT and SA based on their locations with respect to genes, including exonic, intronic, 5’ untranslated region, 3’ untranslated region, upstream, downstream and intergenic, using Ensembl gene definitions.

### Annotating lead SNPs to genes

For each lead SNPs derived from separate conjunctional FDR analysis, we performed positional mapping, eQTL mapping and chromatin interaction mapping in FUMA^50^ using default parameters. These mapping strategies were used to define the functional effects of shared risk loci on the genes. Positional mapping was used to map shared independent lead SNPs to protein-coding genes on the basis of physical distance (within 10 kb) in the human reference assembly (GRCh37/hg19). These genes were then used to examine whether they overrepresented in predefined gene ontology terms (see ‘Gene overrepresentation analysis’ section) and whether they showed differential expression in postmortem brains of individuals with corresponding disorder (see ‘Differential gene expression analysis’ section).

eQTL mapping was used to elucidate the potential functional implications of the identified lead variants. This analysis enabled us to map each lead SNP to genes that exhibit a significant association with their expression levels in brain tissue. Specifically, eQTL mapping was carried out in relation to genes up to 1lJMb away based on four brain-expression data repositories: PsychENCORE^98^, CommonMind Consortium^99^, BRAINEAC^86^ and GTEx v8 Brain^87^. FUMA used an FDR threshold of 0.05 (default parameter) within each analysis to identify significant eQTL associations.

Chromatin interaction mapping was used to map the SNPs to genes exhibiting significant associations with chromatin interaction sites in tissues relevant to the human brain, including: PsychENCORE EP link (one way)^98^; PsychENCORE promoter anchored loops^99^; HiC adult cortex^100^; HiC fetal cortex^100^; HiC (GSE87112) dorsolateral prefrontal cortex^101^; HiC (GSE87112) hippocampus^101^; and HiC (GSE87112) neural progenitor cell^101^. We further selected only those genes for which one or both regions involved in the chromatin interaction overlapped with a predicted enhancer or promoter region (250 bp upstream or 500 bp downstream of the transcription start site) in any of the brain-related repositories from the Roadmap Epigenomics Project^102^. These repositories included: E053 (neurospheres) cortex, E054 (neurospheres) ganglion eminence, E067 (brain) angular gyrus, E068 (brain) anterior caudate, E069 (brain) cingulate gyrus, E070 (brain) germinal matrix, E071 (brain) hippocampus middle, E072 (brain) inferior temporal lobe, E073 (brain) dorsolateral prefrontal cortex, E074 (brain) substantia nigra, E081 (brain) foetal brain male, E082 (brain) foetal brain female, E003 embryonic stem (ES) H1 cells, E008 ES H9 cells, E007 (ES-derived) H1 derived neuronal progenitor cultured cells, E009 (ES-derived) H9 derived neuronal progenitor cultured cells, and E010 (ES-derived) H9 derived neuron cultured cells. An FDR threshold of 1×10^−6^ was applied to identify significant interactions (default parameter), separately for each analysis.

### Multivariate associations with cortical hierarchy maps

A previous study proposed three canonical hierarchies that encompass various large-scale cortical features, including anatomical, functional, and evolutionary hierarchies^54^. The anatomical hierarchy is quantified by the T1-weighted to T2-weighted ratio, with lower hierarchy observed in sensory and motor regions and higher hierarchy in association cortices^103^. The functional hierarchy is measured using the principal gradient of functional connectivity, which aligns spatially with the anatomical hierarchy^57^. The evolutionary hierarchy is reflected by patterns of cortical expansion observed between humans and other species in the mammalian evolutionary tree, with sensory and motor cortical regions dominating the cerebral cortex in early mammals^58^. Building upon these hierarchical frameworks, we sought to investigate whether the cortical regions exhibiting genetic overlap with psychiatric disorders aligned with the cortical organization hierarchy. We used canonical correlation analysis to identify the linear combination of genetic overlap maps of CT and SA with psychiatric disorders that maximally correlated with each cortical hierarchy (https://pennlinc.github.io/S-A_ArchetypalAxis). To assess the significance of the multivariate association analysis for each cortical hierarchy map, spin tests were conducted to randomly spatially rotating the hierarchical ranks relative to the genetic overlap maps in 10,000 iterations. Because cortical data often exhibit distance-dependent spatial autocorrelation, spin tests generate a null distribution by rotating spherical projections of the hierarchical map while maintaining its spatial covariance characterization^104–106^. For significant canonical correlations, we further computed loadings using correlation analysis between the canonical variable and each genetic overlap map across the 180 brain regions. These loadings indicate the extent and direction to which each genetic overlap map contribute to the multivariate association with a specific cortical hierarchy map.

### Gene overrepresentation analysis

Using ConsensusPathDB^60^, we conducted gene overrepresentation analysis to identify shared neurobiological processes associated with the genetic correlates of brain disorders and brain structures. For the conjunctional FDR analysis of each disorder, we integrated all shared risk genes and examined their overrepresentation in predefined gene sets. To perform the analysis, we first defined the background gene set by mapping all SNPs associated with psychiatric disorders in the GWAS to protein-coding genes based on the National Center for Biotechnology Information build 37.3 gene definitions. Each brain disorder had a specific background gene set size, with 18,789 genes for ADHD, 18,886 genes for ANX disorder, 18,873 genes for ASD, 18,943 genes for BD, 18,929 genes for MDD, 18,540 genes for PTSD, and 18,852 genes for SCZ. ConsensusPathDB^60^ utilized the hypergeometric test to assess the overrepresentation of predefined gene sets within the set of shared genes for each brain disorder. This analysis accounted for the size of the background gene set associated with the specific disorder. The tool evaluated the overrepresentation of candidate genes within gene ontology terms, including biological process, molecular function, and cellular component, at four hierarchical levels. For each predefined gene set, the analysis generated a p-value, which underwent multiple testing corrections using a FDR correction threshold of 0.05.

### Differential gene expression analysis

We investigated whether the genes shared between grey matter structures and psychiatric disorders exhibited differential expression in cortical brain tissue obtained from patients with psychiatric disorders compared to healthy controls. To assess this, we used the differential expression profiles from previously published studies of ADHD (24 patients and 29 controls)^66^, ASD (51 patients and 936 controls)^67^, BD (222 patients and 936 controls)^67^, MDD (109 patients and 109 controls)^68^, PTSD (107 patients and 109 controls)^68^, and SCZ (559 patients and 936 controls)^67^ (differential expression data for ANX was not available for comparison^107^). Tissue samples in available studies were obtained from the dorsolateral prefrontal cortex or anterior cingulate cortex. The studies assessed the diagnostic differences in gene expression by deriving *t*-statistics, while adjusting for a set of sensitive confounding factors, such as demographic/clinical features and technical measures. For each disorder dataset, we extracted *t*-values and p-values of differential expression profiles based on genes nearest to the genetic loci shared between brain structures and a given disorder in conjunctional FDR analyses (using p<0.05 threshold to indicate differential expression). Positive and negative *t*-values represented up-regulated and down-regulated expression in patients relative to healthy controls, respectively.

## Acknowledgements

This research was conducted using summary statistics that used the UK Biobank resource under Application Number 20904, with Varun Warrier as the principal applicant. The research was funded by R01MH132934 and R01MH133843.

## Conflict of Interest

AFA-B receives consulting income from Octave Bioscience. AFA-B, JS, and RAIB hold equity in and serve on the board of Centile Bioscience. RTS receives consulting income from Octave Bioscience and compensation for scientific reviewing from the American Medical Association. OAA is a consultant to cortechs.ai, and has received speaker’s honorarium from Lundbeck, Janssen, Sunovion.

